# Multi-trait genomic prediction improves selection accuracy for enhancing seed mineral concentrations in pea (*Pisum sativum L.*)

**DOI:** 10.1101/2022.04.11.487944

**Authors:** Sikiru Adeniyi Atanda, Jenna Steffes, Yang Lan, Md Abdullah Al Bari, Jeonghwa Kim, Mario Morales, Josephine Johnson, Rica Amor Saludares, Hannah Worral, Lisa Piche, Andrew Ross, Michael A Grusak, Clarice J. Coyne, Rebecca J. McGee, Jiajia Rao, Nonoy Bandillo

## Abstract

The superiority of multi-trait genomic selection (MT-GS) over univariate genomic selection (UNI-GS) can be improved by redesigning the phenotyping strategy. In this study, we used about 300 advanced breeding lines from North Dakota State University (NDSU) pulse breeding program and about 200 USDA accessions evaluated for ten nutritional traits to assess the efficiency of sparse testing in MT-GS. Our results showed that sparse phenotyping using MT-GS consistently outperformed UNI-GS when compared to partially balanced phenotyping using MT-GS. This strategy can be further extended to multi-environment multi-trait GS to improve prediction performance and reduce the cost of phenotyping and time-consuming data collection process. Given that MT-GS relies on borrowing information from genetically correlated traits and relatives, consideration should be given to trait combinations in the training and prediction sets to improve model parameters estimate and ultimately prediction performance. Our results point to heritability and genetic correlation between traits as possible parameters to achieve this objective.

## 1.0 Introduction

In recent times, there is an increased demand for genetic improvement of nutritional traits in crops due to the growing demand for plant-based protein, mineral elements and vitamins. Pulse crops are known to have high protein value and are rich in micronutrients with potential to alleviate hidden hunger (Mudryj et al. 2014; Wadhawan et al. 2021; Bari et al. 2021). However, phenotyping/screening for nutritional traits such as protein, manganese, selenium, copper, zinc, iron, potassium, phosphorus, magnesium and calcium is expensive and time consuming, especially in the early yield testing stage with hundreds of lines to evaluate. This is a major limitation in a public breeding program aiming to have a biofortified product profile. However, due to advancement in the genotyping platform, the cost of genotyping is becoming relatively less expensive compared to the cost of phenotyping; thus, genomic selection (GS) that uses whole-genome information to predict genomic estimated breeding value (GEBV) of unobserved genotypes is gaining traction as breeders’ choice of selection method (Poland et al. 2012; Zhao et al. 2021; Bassi et al. 2016; Santantonio et al. 2020; Atanda, et al. 2021a). Though GS research in pea is scanty, the available studies (Annicchiarico et al. 2019; Crosta et al. 2021; Bari et al. 2021) show GS potential to predict the genetic merit of pea lines and germplasm accessions. Following Bari et al. (2021), the North Dakota State University (NDSU) pulse breeding program is prioritizing the use of GS particularly in the preliminary yield trial (PYT or stage 1) where effectiveness of phenotypic selection is limited by phenotyping in one/two locations due to seed multiplication challenges for hundreds of lines for multi-location trials. Consequently, the NDSU pulse breeding program is focused on redesigning the PYT from phenotypic based selection to GS to reduce the number of seeds for phenotyping and increase selection accuracy for advancement of promising lines to advanced yield testing stage.

In general, GS is often performed with univariate-trait (UT) models that assume genetic correlation among traits to be zero (Jia and Jannink 2012; Montesinos-López et al. 2016, 2018; Bhatta et al. 2020; Gaire et al. 2022). However, in practice, breeders’ select for multiple traits that are genetically correlated, ranging from negative to positive correlations. To harness the genetic correlation between traits and among genotypes to improve prediction accuracy, multi-trait (MT) models, which are the generalization of UT models, have been investigated. Several empirical studies (Calus and Veerkamp 2011; Montesinos-López et al. 2018; Bhatta et al. 2020; Gaire et al. 2022) have reported improved prediction accuracy in different crops using MT models that allow borrowing of information between correlated traits and among genotypes compared to UT models. Prediction accuracy in MT-GS improves as correlation between traits increases (Jia and Jannink 2012; Okeke et al. 2017; Montesinos-López et al. 2018, 2019; Neyhart et al. 2019), however, trait heritability varies and is a key limiting factor to upper bound of prediction accuracy (Manolio et al. 2009; Yang et al. 2015; Schopp et al. 2017; Zhang et al. 2019; Atanda et al. 2021a). These factors will likely influence the composition of traits in the training and the prediction sets in MT-GS and ultimately the prediction accuracy.

In the MT-GS model, the training set consists of individuals with phenotypic records for all traits to predict the genetic values of un-phenotyped individuals in the prediction set using genome-wide marker information. The crucial question is how to design a MT-GS strategy that will optimize the trade-off between the limiting factors and accuracy of predicting the genetic value of the traits. Studies (Montesinos-López et al. 2016, 2018, 2019; Guo et al. 2014; Bhatta et al. 2020; Gaire et al. 2022) have highlighted the importance of each factor to prediction accuracy; however, nothing is known about their combinations on composition of traits in the training and prediction set in the context of MT-GS. Consequently, we investigated the influence of the limiting factors on composition of traits in the training and prediction sets to guide the use of MT-GS in a breeding program.

Further, in most MT-GS cross-validation studies (Montesinos-López et al. 2016, 2018, 2019; Guo et al. 2014; Bhatta et al. 2020; Gaire et al. 2022), the same set of genotypes overlap across traits for testing prediction models (**Suppl. Fig. 1A, C**). In such a scenario, the same set of genotypes have phenotypic records for all traits while the other genotypes serve as a prediction set (partially balanced testing strategy, denoted as PBT); however, results from multi-environment GS studies have shown that this approach is less optimal compared to sparse testing using genomic prediction where phenotyping of genotypes is split across environments (Burgueño et al. 2012; Jarquin et al. 2020; Atanda et al. 2021). We extend sparse phenotyping in the context of MT-GS in which the phenotyping of lines is split across traits (**Suppl. Fig. 1B, D**). This strategy could improve prediction accuracy in MT-GS by efficiently using information across traits and genotypes. More so, it can be robust for building historical data for use in prediction models, since all genotypes have phenotypic records for the different traits. To further evaluate the potential of GS in the NDSU pulse breeding program and how it can be efficiently deployed to improve genetic gain, the following were our objectives in this study: 1) determine the efficiency of MT compared to UT in predicting nutritional traits in pea, 2) determine the optimal method to design training and testing trait sets using heritability and genetic correlation between traits as metric, and 3) identify optimal resource allocation for phenotyping nutritional traits in the early yield testing stage by comparing the predictive ability of sparse and partial balanced testing.

## 2.0 Materials and Methods

### 2.1 Genetic Materials and Field or Greenhouse Evaluation

The genetic material consisted of 282 pea lines (DS1) from North Dakota State University (NDSU) pulse breeding program and 192 USDA pea accessions (DS2) previously described in Bari et al. (2021). The NDSU lines were planted in augmented row-column design with five repeated checks in the 2020-2021 growing season at the North Dakota Agricultural Experiment Station, Minot, North Dakota, United States (27°29′N, 109°56′W). Seeds were treated with fungicide and insecticide prior to planting. At planting, 30 seeds were planted on 152 × 60 cm plot size with 30 cm spacing between plots. Plots were harvested at physiological maturity (90-120 days after planting) and dried to 15% moisture content. For the USDA pea accessions, six plants of each accession were grown in 5L black plastic pots filled with a synthetic soil mix composed of two parts Metro-Mix 360 (Scotts-Sierra Horticultural Products Co., Marysville, Ohio) and one part vermiculite (Strong-Lite Medium Vermiculite, Sun Gro Horticulture Co, Seneca Illinois). Plants were grown in a controlled environment greenhouse with a temperature regime of 22 ± 3 °C/day and 20 ± 3 °C/ night, with a relative humidity ranging from 45% to 65% throughout the day/night cycle. Sunlight was supplemented with metal halide lamps, set to a 15 h day, 9 h night cycle (lights on at 700 h). In order to maintain an adequate supply of all mineral nutrients, a complete fertilizer mixture was provided to each pot on a daily basis. Pots were irrigated with an automated drip irrigation system (one drip line to each pot); the system was regulated with a timer that delivered nutrient solution twice a day (younger plants) or three times a day (older plants) in sufficient quantity to saturate the soil mass at each irrigation. The nutrient solution contained the following concentrations of mineral salts: 1.0 mM KNO_3_, 0.4 mM Ca(NO_3_)_2_, 0.1 mM MgSO_4_, 0.15 mM KH_2_PO_4_ and 25 μM CaCl_2_, 25 μM H_3_BO_3_, 2 μM MnSO_4_, 2 μM ZnSO_4_, 0.5 μM CuSO_4_, 0.5 μM H_2_MoO_4_, 0.1 μM NiSO_4_, 1 μM Fe(Ш)-N, N’-ethylenebis[2-(2-hydroxyphenyl)-glycine] (Sprint 138; Becker-Underwood, Inc., Ames, Iowa, USA). We thus attempted to maintain all essential minerals at sufficient, but non-toxic levels, in the soil. Seeds were harvested from each accession at full plant maturity.

### 2.2 Mineral Analysis

Mineral elements for DS1 were measured following procedures described in (Ma et al., 2017; Lan et al. 2019). Briefly, 200g non-dehulled pea seeds were ground to fine flour and then digested with concentrated nitric acid (70% HNO_3_) in a digestion system block at 90 °C for 60 minutes. Afterward, 3 mL of hydrogen peroxide was added to further the digestion process for 15 minutes, followed by the addition of 3 mL hydrochloric acid (70% HCl) and heated for additional 5 minutes. After cooling to room temperature, the digested samples were filtered through DigiFILTER (SCP Science) and diluted to 10mL with nanopure water. To validate the procedure and analytical measurement, an apple leaf standard (SRM 1515; National Institute of Standards and Technology, Gaithersburg, Maryland, USA) was analyzed simultaneously with the pea flour samples. Total concentration of the mineral elements was measured using inductively coupled plasma atomic emission spectrometry (IRIS Advantage ICP-AES; Thermo Elemental, Franklin, Massachusetts, USA). Mineral values were determined with the ICP-AES using the following spectral emission lines (in nm): Ca, 184.0; Mg, 285.2; K, 769.8; P, 177.4; Fe, 238.2; Zn, 213.8; Mn, 260.5; Cu, 324.7; Ni, 231.6; B, 208.9; Mo, 202.0.

For the DS2, dried seeds (with seed coats) from 6 plants of each accession were ground to a uniform powder using a coffee grinder with stainless steel blades. Two sub-samples of each accession were weighed (approximately 200 mg each), dry ashed, resuspended in ultra-pure nitric acid and analyzed for Ca, Mg, K, P, Fe, Zn, Mn, Cu, Ni, B and Mo concentrations using inductively coupled plasma atomic emission spectrometry (IRIS Advantage ICP-AES; Thermo Elemental, Franklin, Massachusetts, USA). Dry ashing was performed in quartz tubes, with samples ashed for 6 h at 450 °C. After cooling, to ensure complete oxidation of all tissues, 2.5 ml of 30% H_2_O_2_ was added to each tube and samples were reheated to 450 °C for 1 h. Apple leaf standards (SRM 1515; National Institute of Standards and Technology, Gaithersburg, Maryland, USA) were ashed and analyzed along with pea seed samples to verify the reliability of the procedures and analytical measurements. Mineral values were determined with the ICP-AES using the same spectral emission lines (in nm) noted for the DS1 population.

### 2.3 Genotyping

Details on DNA isolation and genotyping-by-sequencing (GBS) can be found in Bari et al. (2021). DS1 and DS2 were genotyped using GBS and 28, 832SNP markers were generated for DS1 while 380,527 SNP markers were generated for DS2. After removing SNPs with more than 90% missing values, heterozygosity greater than 20% and with a minor allele frequency less than 5%, 11, 858 and 30, 645 SNPs remained for DS1 and DS2 respectively and were used for the analysis. Missing SNPs were imputed with Beagle 5.1 (Browning et al., 2018).

### 2.4 Phenotypic Data Analysis

Best linear unbiased estimates of the phenotypes for DS1 accounting for spatial trend on the field modelled by a smooth bivariate function of the spatial coordinates f(**r, c**) represented by 2D P-splines was implemented in SpATS R package (Rodríguez-Álvarez et al. 2016). This was modeled as:

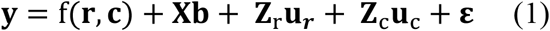

where: **y** is the response variable for n-th phenotype, **b** is the fixed effect of the genotype, **u**_**r**_ and **u**_**c**_ are row and column random effects accounting for discontinuous field variation with multivariate normal distribution: 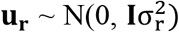 and 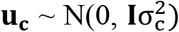 respectively. **I** is an identity matrix and 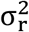 and 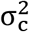 are variance for row and column effect. f(**r, c**) is a smooth bivariate function defined over the row and column positions (see Velazco et al. 2017 for details), **ε** is the measurement error from each plot with distribution of 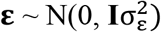. **I** is the same as above, 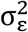 is variance for the residual term or simply referred to as nugget. **X** and **Z** are incidence matrix for the fixed and random terms.

For the DS2, the mineral elements value of each accession was estimated as follows; mineral values from the two sub-samples (see Mineral Analysis section for details) were averaged for each accession; these averaged values are presented as ppm (parts per million), which is equivalent to ug/g DW (micrograms per gram dry weight). In this study it was denoted as mean phenotypic value of each accession for each mineral element. In general, the standard deviations for each mineral were low (i.e., within each accession). Across all accessions, the average standard deviation for each mineral (calculated as percent of the mean of the two sub-samples) was: Ca, 10.4%; Mg, 2.0%; K, 2.5%; P, 2.5%; Fe, 6.0%; Zn, 5.1%; Mn, 6.9%; Cu, 7.5%; Ni, 15.4%; B, 5.8%; Mo, 3.7%.

### 2.5 Genomic Selection Models

Univariate GS model was implemented in ‘BGLR’ R package (Pérez and de los Campos 2014) using Bayesian ridge regression model equivalent to the genomic best linear unbiased prediction (GBLUP) model expressed as:

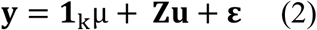

where y is the vector (n × 1) of adjusted means (BLUEs) using DS1 or mean phenotypic value using DS2 for k-th genotypes for a given n-th trait/mineral element, μ is the overall mean and **1**_k_ (kx1) is a of vector ones, **u** is the genomic effect of k-th genotypes assumed to follow multivariate normal distribution expressed as 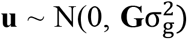. **G** is the genomic relationship matrix and 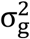 is the additive genetic variance.

MT-GS model was fit using Bayesian multivariate gaussian model in ‘MTM’ R package (de los Campos and Grüneberg 2016). This is expressed as:

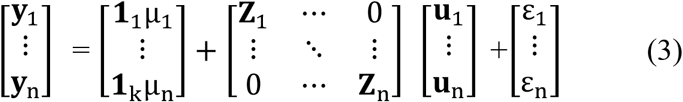

where **y**_1_…**y**_n_ are the vector of phenotypes, μ_1_ …μ_n_ are the overall mean for each n-th trait, **Z**_1_ …**Z**_n_ is the incidence matrix for genomic effect of the lines for each n-th trait, **u**_1_ … **u**_n_ is genomic effect of the lines for each n-th trait and **ε**_1_ … **ε**_n_ is the residual error for each n-th trait. The random term is assumed to follow multivariate normal distribution [**u**_1_ … **u**_n_] ∼ N[0, (**G ⨂ G**_o_)]. Where **G** is the same as above and **G**_0_ is an unstructured variance-covariance matrix of the genetic effect of the traits, this is represented as follows:

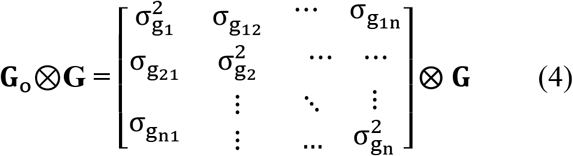

The off-diagonal elements represent variance for each trait and covariances between traits are the off-diagonal elements.

Further, the residual term for each n-th trait is assumed to follow multivariate normal distribution: [**ε**_1_ … **ε**_n_] ∼ N[0, (**I ⨂ R**)], where **I** is the same as above and **R** is a heterogeneous diagonal matrix of the residual variances for each n-th trait:

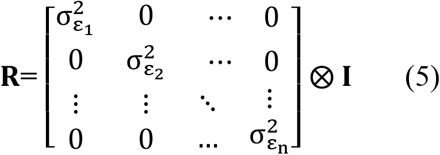

The diagonal elements represent the residual variance for each n-th trait and off-diagonal elements of the **R** matrix equal zero. In our preliminary analysis unstructured **R** matrix where off-diagonal element of **R** represent covariance of the residual effects of the traits was considered; however, we observed inconsistent model convergence for all iterations. The same results were observed when factor analytic model was considered for the **R** structure which might be due to size of the dataset used in our study relative to the number of model parameters to estimate.

Genomic heritability estimate (de los Campos et al. 2015; Feldmann et al. 2021) for n-th trait using individual level data was derived from the variance components obtained from the model using the complete dataset.

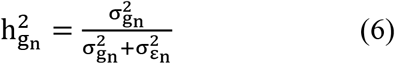

where 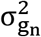 and 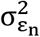 are the genetic, residual variance estimates for n-th trait.

### 2.4 Cross-Validation Scheme

To evaluate the performance of sparse testing strategy in the context of MT-GS, different cross-validations mimicking potential applications of MT-GS in a breeding program were explored. Leveraging on the results from sparse testing in multi-environment yield trials using GS (Jarquin et al. 2020; Atanda et al. 2021b), we varied the number of genotypes that serve as connectivity across traits to assess predictive ability in the different scenarios. Depending on the size of the data set and the number of phenotypes, different overlapping sizes were evaluated (**Suppl. Table 1**). Five different overlapping sizes (50, 60, 70, 80, 90%) were considered for DS1 (n=282), which had the highest total number of genotypes, followed by four overlapping sizes (40, 50, 60, 70%) in DS2 (n=192). For example, when 50% of the total genotypes in DS1 serve as connectivity across the traits, the remaining 141 genotypes were partitioned into 10 distinct sets, each trait with a unique set. Thus, each trait has 155 genotypes as training set to predict the genetic merit of 127 genotypes (**Suppl. Figure 1B**). This process was repeated 50 times. As the size of the overlapping genotypes increased (60, 70, 80, and 90% of total genotypes), the training set size increased to 180, 205, 230, 255, and the prediction set size reduced to 102, 77, 27 respectively. The splitting of the genotypes across traits was also repeated 50 times for each overlapping size scenario, each iteration has different genotypes that serve as connectivity across traits, non-overlapping training set for each trait and the prediction set (**Suppl. Table 1**). In each iteration, the Pearson correlation of the predicted GEBV and the BLUE estimates of the genotypes for each trait obtained using complete dataset was calculated and the mean was recorded as the predictive ability of the prediction set for each trait. The same process was repeated for DS2, however, the predictive ability in each iteration is the Pearson correlation of the predicted GEBV obtained using complete dataset and the mean phenotypic value of each accession for each trait and the mean was recorded.

To determine the efficiency of MT (sparse and partially balanced phenotyping) and univariate (UNI) GS model, we compared the predictive ability of the prediction sets using the different training set size defined in each dataset. Again, this process was repeated 50 times, each iteration having different genotypes included in the training and prediction set for all traits in the UNI-GS model and across traits for the partially balanced phenotyping MT-GS. For sparse phenotyping each iteration has different genotypes that serve as connectivity across traits, a non-overlapping training set for each trait, and the prediction set (**Suppl. Figure 1**). For DS1, predictive ability for each iteration was measured as the Pearson correlation of the predicted GEBV and the BLUE estimates of the genotypes for each trait obtained using the full dataset. Average was reported. In the DS2, BLUE was replaced with the mean phenotypic value of each accession for each trait.

Based on the preliminary analysis results, only the sparse testing using MT-GS was considered to evaluate the efficiency of using heritability, genetic correlation between traits or combination of the factors for trait assignment in the prediction set and/or training set respectively. The following scenarios were assessed:

1. Exclusion of trait(s) with lowest heritability but moderate to high genetic correlation with other traits from the prediction set however reserved in the model. We also evaluated the scenario when it was removed from the model.
2. Exclusion of trait(s) with the highest occurrence of negative correlation with other traits but moderate to high heritability from the prediction set but reserved in the model. We also evaluated the scenario when it was removed from the model.
3. Exclusion of trait(s) with the lowest heritability but moderate to high genetic correlation with other traits, as well as trait(s) with the highest occurrence of negative correlation with other traits but moderate to high heritability, from the prediction set but reserved in the model. We also evaluated the scenario in which they were left out of the model.

## 3.0 Results

### 3.1 Traits genomic heritability (diagonal) and genetic correlation among traits (upper diagonal)

In DS1 heritability was moderately high for all traits except Ca with very low heritability value of 0.01 (**Fig. 1A**). However, in DS2 Ca had moderate heritability of 0.40 (**Fig. 1B**). Similarly, Fe had heritability of 0.63 in DS2 compared to 0.29 in DS1, and in general the traits heritability in DS2 ranged from moderate to high. In the two datasets, P consistently had the highest heritability of 0.87 in DS1 and 0.73 in DS2. The genetic correlation between traits in DS1 ranged from -0.01 to 0.96 while it ranged from -0.01 to 0.99 in DS2. In DS1, Cd had zero or no genetic correlation with most of the traits. Similarly, K had no genetic correlation with Ca in DS2, contrary to the 0.33 genetic correlation observed in DS1 (**Fig. 1A, B**). Generally, in DS1, K, Fe, P and Mg had moderate to high genetic correlation with most of the traits while in DS2 Ni, Cu and Mg had high genetic correlation with other traits except with Mo.

**Figure 1:**
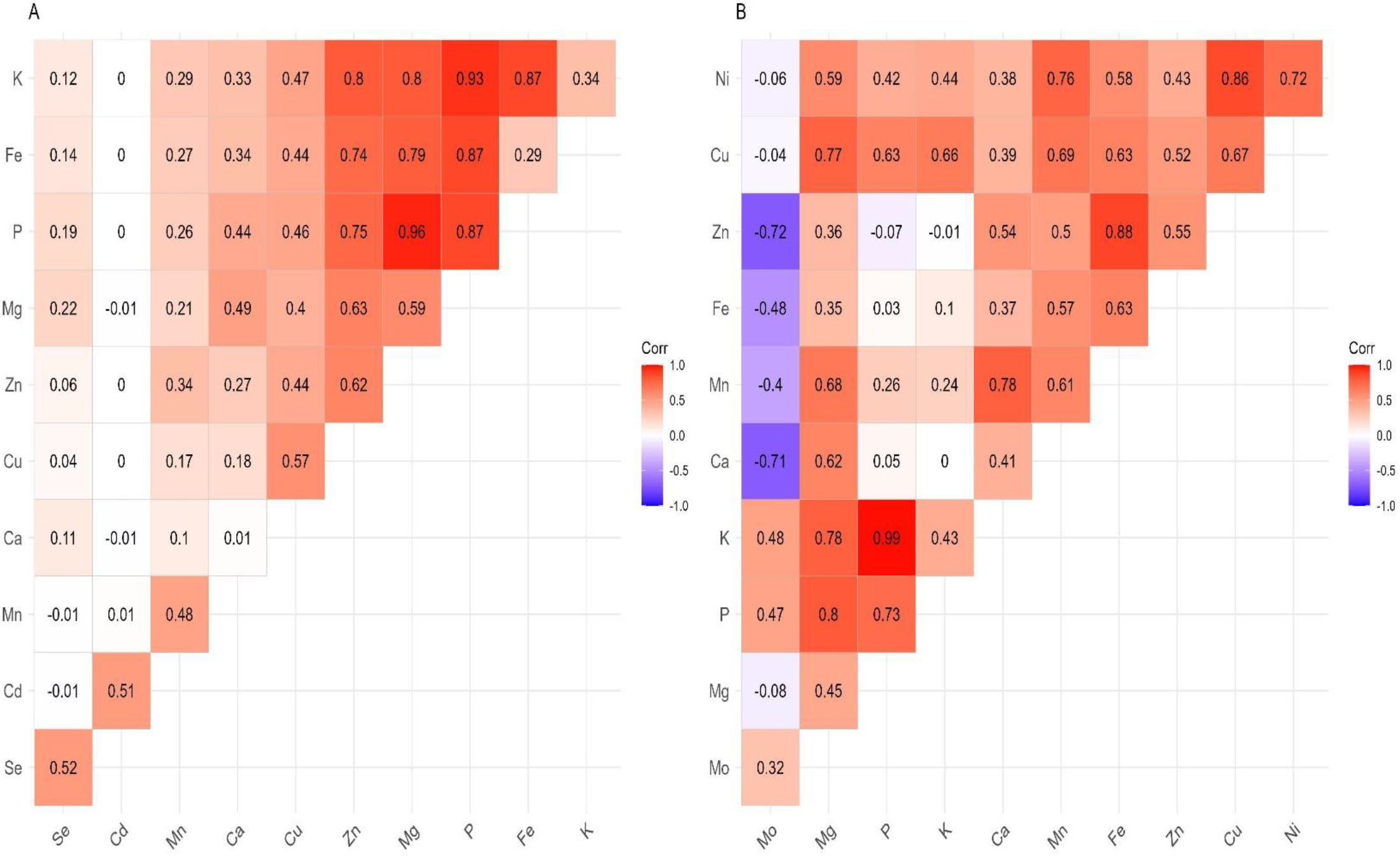
Genomic heritability (diagonal) and genetic correlation between pairs of traits (upper diagonal) from MT-GS model using complete datasets. Fig. 1A, represent results using DS1 dataset and Fig. 1B indicate results using DS2 dataset.

### 3.2 Sparse testing MT-GS improves predictive ability across traits compared to partially balanced testing MT and univariate GS models

Regardless of cross-validation schemes or dataset, sparse testing MT-GS model outperformed PBT MT and UNI-GS models for all traits except for Ca in DS1 which might be attributed to near-zero genetic signal observed for this trait (**Fig. 2A: B**). For instance, in DS2 where the predictive ability is generally high compared to DS1, sparse testing using MT-GS outperformed PBT using MT-GS by 25, 36, 15, 26, 27, 67, 81, 50, 66, 56% respectively for Mo, Mg, P, K, Ca, Mn, Fe, Zn, Cu and Ni while it improved predictive ability by 7, 58, 23, 17, 60, 12, 2, 95, 56% compared to UN-GS model (**Fig. 2B**). Surprisingly, PBT using MT-GS model did not consistently outperform UNI-GS model in DS2 compared to DS1 where PBT using MT-GS marginally results in improved predictive ability for all traits.

**Figure 2:**
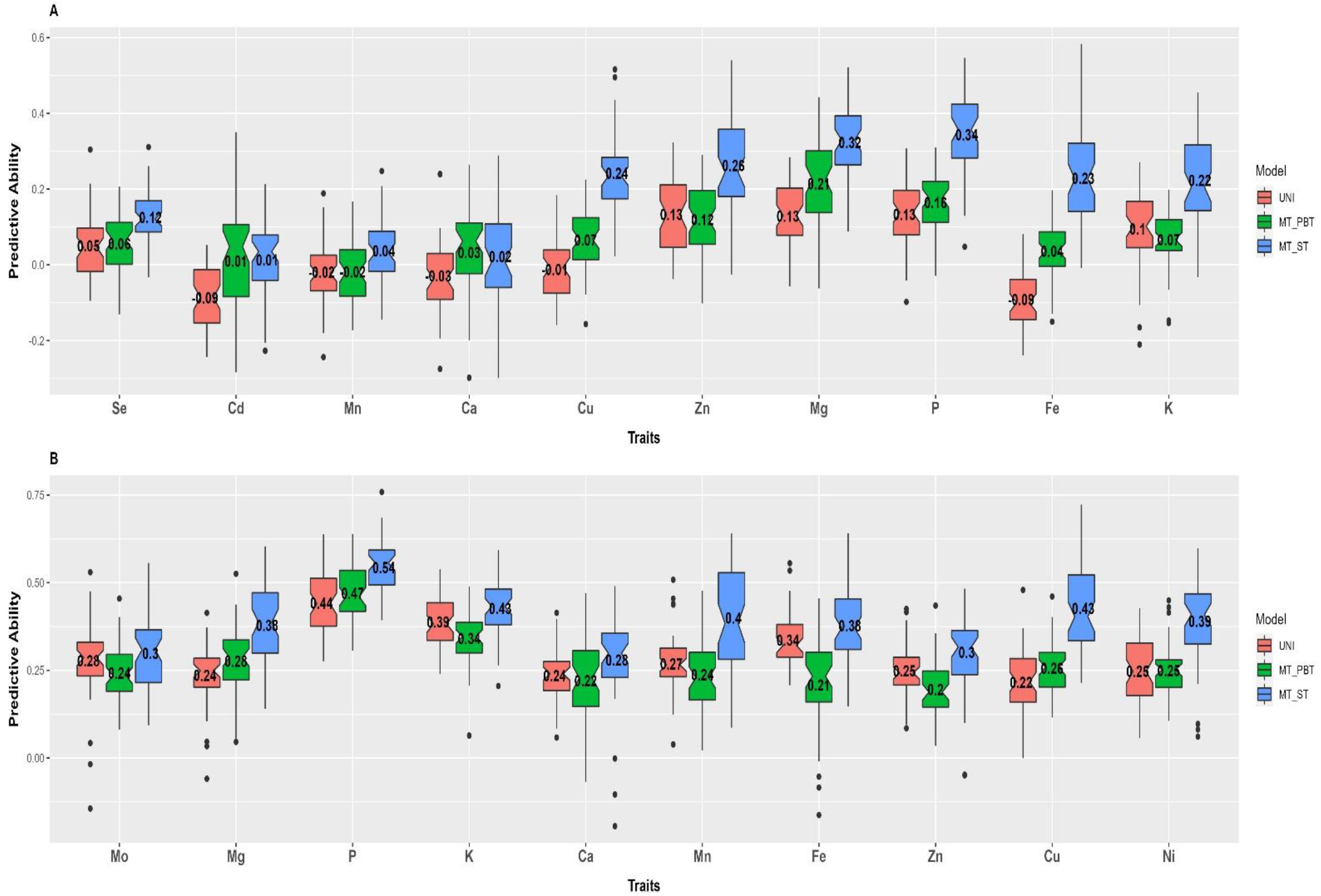
Predictive performance of UNI and MT-GS using partially balanced (PBT) and sparse testing (ST) phenotyping of the traits. The number within each box represents mean predictive ability of 50 iterations of the process of line assignment as training and prediction set. In each iteration different genotypes were assigned as training and prediction set for the traits in the UNI-GS model and across traits for the partially balanced phenotyping MT-GS. For sparse phenotyping each iteration has different genotypes that serve as connectivity across traits, a non-overlapping training set for each trait, and the prediction set

### 3.2 Traits combination as a function of heritability, genetic correlation between traits and their combination

When either heritability or genetic correlation was considered a decision tool for a combination of traits in the prediction and/or calibration set, the predictive ability improved for all traits compared to having all the traits in the prediction and the training set. However, the magnitude of the gain in predictive ability varies by trait (**Fig. 3, 4**). In DS1, for example, when Ca with the lowest heritability (0.01) but moderate to high genetic correlation with other traits was dropped from the prediction set but kept in the model, the gain in the predictive ability for the remaining 9 traits in the prediction set across the overlapping scenarios ranged from 2.79 to 85.07% (**Fig. 3B**), while it ranged from 3.28 to 63.37% when excluded from the model (**Fig. 3B\***). When Cd with high heritability (0.51) but zero genetic correlation with most traits was removed from the prediction set, but retained in the calibration model, the improvement in predictive ability ranged from 6.75 to 104.41% (**Fig. 3C)** and ranged from 1.38 to 45.42% when removed from the training model (**Fig. 3C*)**. Similar results were obtained in DS2, when Mo with heritability of 0.43 and negative correlation with the majority of the traits was removed from the prediction set but reserved in the calibration model. The predictive ability of traits in the prediction set ranged from 2.41 to 77.92% (**Fig. 4B)** and ranged from 0.62 to 19.62% when removed from the model (**Fig. 4B*)**. Because Mo has a negative genetic correlation with the majority of the traits, in addition to having the lowest heritability of all the traits in DS2, we substitute Mo with Ca, which has a heritability of 0.41 and a moderate to high genetic correlation with other traits, to disentangle the confounding effect of heritability and genetic correlation. The gain in predictive ability ranged from 3.19 to 90.34% when the calibration model was reserved (**Fig. 4C**) and from 1.34 to 14.65% when the calibration model was removed (**Fig. 4C\***).

**Figure 3:**
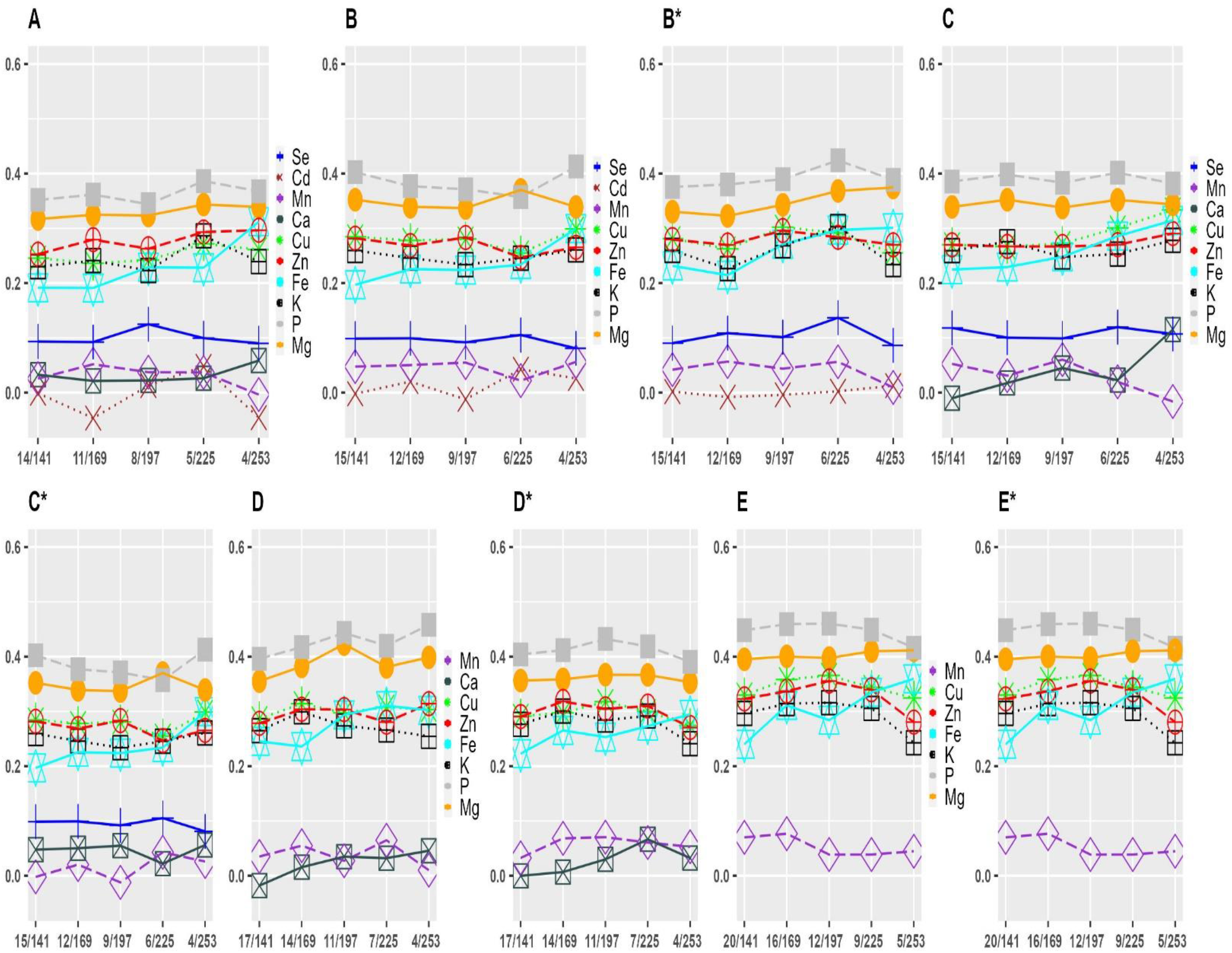
Predictive ability of untested lines in DS1 for each trait for different overlapping and non-overlapping size. The different colors denote traits in the prediction set, which might also be present in the calibration model. The suffix (*) indicate exclusion of trait (s) from the calibration model based on its heritability, degree of genetic correlation with other traits or combination of the two factors.

**Figure 4:**
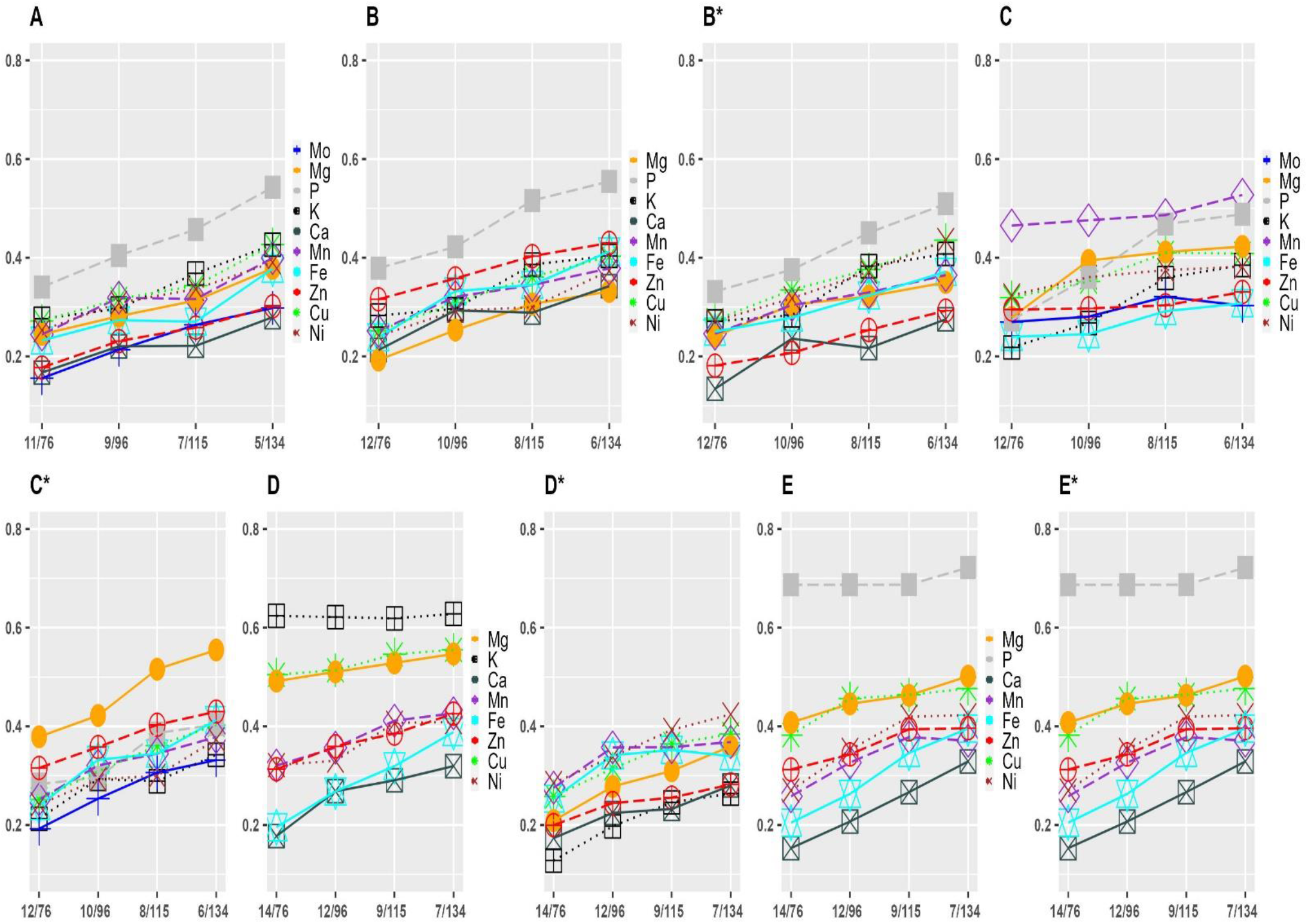
Predictive ability of untested lines in DS2 for each trait for different overlapping and non-overlapping size. The different colors denote traits in the prediction set, which might also be present in the calibration model. The suffix (*) indicate exclusion of trait (s) from the calibration model based on its heritability, degree of genetic correlation with other traits or combination of the two factors.

Unsurprisingly, when Mo and P, which have moderate and high heritabilities of 0.32 and 0.73 but are negatively correlated with other traits, respectively, were excluded from the prediction set, the predictive ability improved for all traits except Ca and Fe (**Fig. 4D**). When the traits were removed from the calibration model, the predictive ability decreased for majority of the traits (**Fig. 4D\***). On the contrary, removing Mo and K from the prediction set with moderate heritability of 0.32 and 0.43 resulted in an improved predictive ability (**Fig. 4E, E\***). In contrast to P, K has zero, weak negative, and strong positive correlations with other traits. Figures 3D and 3D* corroborate the findings in Figures 4D and 4D*, in which Se and Cd with moderate heritability of 0.51 and 0.52, respectively, but low genetic correlation with other traits, were excluded from the prediction set but retained or removed from the calibration model. The additional improvement in predictive ability observed when Se, Cd, and Ca were excluded from the prediction set (**Fig. 3E, E\***) demonstrates the efficacy of heritability and genetic correlation between traits as decision metric, corroborating the results obtained in Figure 4E and 4E*.

## 4.0 Discussion

Breeders make advancement decisions based on multiple traits with varying genetic correlations, ranging from negative to positive, and in exceptional cases, no genetic correlation at all. Thus, the use of the MT-GS model is gaining popularity as a choice GS model to estimate the genetic merit of new genotypes. When comparing models, our results corroborate previous studies (Calus and Veerkamp 2011; Jia and Jannink 2012; Montesinos-López et al. 2018; Lado et al. 2018; Bhatta et al. 2020; Gaire et al. 2022) that MT-GS outperforms UNI-GS by harnessing genetic correlation between traits to improve predictive ability across traits. The proposed MT-GS aided sparse phenotyping depart from the previous reports of weak genetic correlation between traits as a limitation to the advantage of MT-GS over UNI, which was evident in the partially balanced phenotyping aided MT-GS. The performance of MT-GS enabled sparse phenotyping was consistently superior to UNI-GS, with least 20% improvement on predictive ability on average across traits, suggesting the importance of borrowing information across traits and related genotypes. Similar results have been reported in sparse testing aided GS in multi-environment trials (Atanda et al. 2021a; 2021b; 2022). This demonstrates the improvement in predictive performance in sparse testing using MT-GS is primarily due to efficient estimation of correlated effects across genetically related traits, as phenotypic records are available for all traits, albeit in a different set of genotypes. In addition, allowing for significant genotype overlap improves predictive ability because genetic connectivity across traits improves estimates of trait-to-trait correlation effects. The observed inflection points in this study, however, suggests that more research is required to determine the optimal number of overlapping genotypes, which might be influenced by the degree of genetic relationship between lines, the number of lines per cross, the genetic correlation between traits, yield testing stage and expected prediction accuracy. Overall, predictive ability improves with heritability in all models except Se, Cd, and Mn in DS1, though DS1 generally has low predictive ability compared to DS2, presumably due to low genetic variation for nutritional traits in DS1 which are elite breeding lines compared to DS2 which are accessions and the growing condition of the accessions in the greenhouse compared to DS1 planted out in the field. Thavarajah et al. (2022) reported heritability estimates of nearly zero for Ca, K, P, Mg, Mn, Fe, Zn, Cu and Se in 44 pea lines evaluated in two locations in 2019 and one location in 2020 with two replications in each location. On the contrary, Ma et al. (2017) observed moderately high genetic diversity for mineral elements in 158 recombinant inbred lines evaluated in two locations with two replications. Given the number of replications and locations used in these studies further suggests that the degree of genetic variation (by inference heritability) for nutritional traits in DS1 may be responsible for the observed low predictive ability.

Multi-trait combinations were created in training and prediction sets based on genetic correlations between traits, heritability, and the combination of the limiting factors to optimize the trade-off between the limiting factors and the accuracy of predicting the genetic value of the phenotypes. In general, the improvement in predictive ability when traits with low heritability but moderate to high genetic correlation with other traits or traits with high occurrence of negative correlation with other traits but moderate to high heritability were excluded from the prediction set suggests that traits with low heritability or genetic correlation with other traits cannot be adequately predicted (Jia and Jannink 2012; Gaire et al. 2022). However, reserving the traits in the training set as secondary traits improves estimation of model parameters resulting in an improvement in predictive ability compared to exclusion from the model. The observed difference in predictive ability for each limiting factor suggests both factors independently affects predictive ability. Consequently, both factors are equally important in determining traits combination in MT-GS. In practice, this information can be sourced from relevant literature on the phenotypes or historical data in the breeding program. To our knowledge this is the first time these two factors are designed to designate traits in the training and prediction set to improve predictive performance in MT-GS. The gain in predictive performance achieved by using this strategy requires further investigation because it has only been tested in pea datasets with limited environments (year by location combinations) and replication which is a major limitation in this study, and does not represent extensive data generated in breeding programs. The availability of multi-environment dataset can improve estimate of genotypic values for quantitative traits. Since significant progress has been made in multi-trait multi-environment genomic prediction (Montesinos-López et al. 2016, 2018, 2019; Gill et al. 2021; Sandhu et al. 2022), our findings suggest future research should focus on developing an optimal strategy for genomic prediction enabled sparse testing of multiple traits in multi-environment trials. This will likely further lower the cost of phenotyping and the time-consuming data collection process. In addition, we encourage use of different crops with varying genetic backgrounds that fairly cover the diversity of data generated in breeding programs to gather more evidence on the efficiency of this strategy in improving prediction performance in MT-GS.

## Conclusion

In this study we propose use of sparse testing in MT-GS which ultimately can be extended to multi-environment multi-trait GS to improve prediction performance and further reduce the cost of phenotyping and time-consuming data collection process. Although our results agree with previous study that weak correlation is a limiting factor of MT-GS superiority over UNI-GS using partially balanced phenotyping. However, our results were inconsistent with sparse phenotyping, suggesting that MT-GS performance can be improved further if phenotyping strategy is redesigned. Our results show that traits combination in training and prediction sets impact prediction performance. Therefore, when designing MT-GS strategy, consideration should be given to traits combination in the training and prediction sets. In addition, our results suggest the use of heritability and genetic correlation between traits as metrics to achieve this objective.

## Data Availability Statement

The SNP dataset used in this study is available via: https://www.ncbi.nlm.nih.gov/sra/PRJNA730349. And the phenotypic data will be made available by reaching out to the corresponding author.

## Author Contributions

SA and NB conceptualized the study. SA performed the analyses, and wrote the manuscript. HW, AR, MM, JK, and MAB coordinated the field experiments for the NDSU breeding lines. LP, JS, JJ and RAS coordinated the genotyping experiments for the NDSU breeding lines. JR, YL, JS, and LP coordinated the mineral profiling of the NDSU breeding lines. MAG, CJC, and RJM designed and executed the greenhouse experiment, genotyping, and mineral profiling of the USDA accessions. NB oversaw the statistical analyses and contributed to the writing of the manuscript. All authors edited, reviewed, and approved the manuscript.

## Acknowledgments

This research was supported through funding from USDA Plant Genetic Resource Evaluation, USA Dry Pea and Lentil Council Research Committee, USDA ARS Pulse Crop Health Initiative and USDA ARS Project: 5348-21000-017-00D (CC), and 5348-21000-024-00D (RJM). Also, grants from USDA-NIFA (Hatch Project ND01513 for NB) and the North Dakota Department of Agriculture through the Specialty Crop Block Grant Program (19–429).

---

**Supplementary Table 1:**
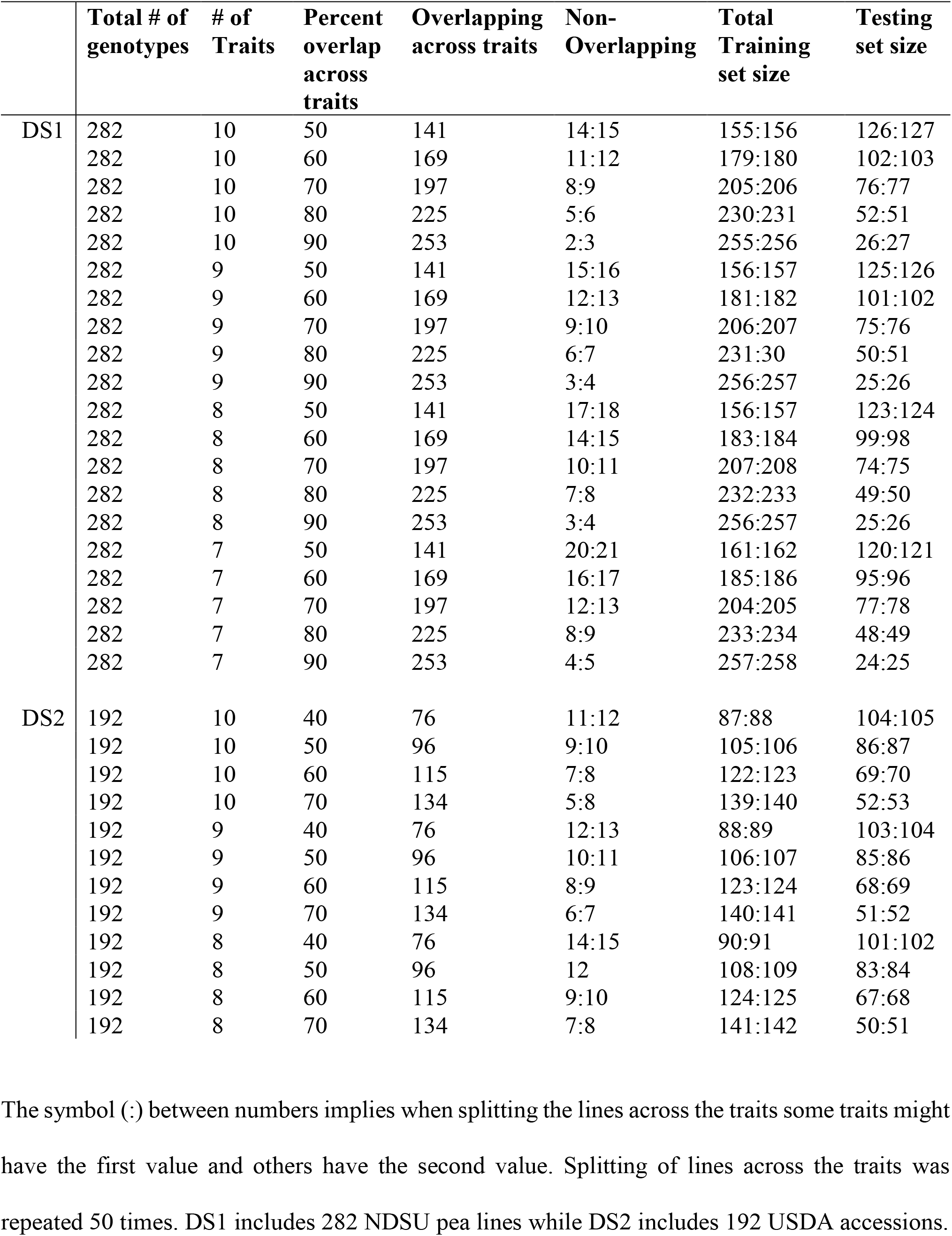
Proportion of genotypes overlapping and non-overlapping across traits for DS1 and DS2 datasets.

|  | Total # of<br>genotypes | # of<br>Traits | Percent<br>overlap<br>across<br>traits | Overlapping<br>across traits | Non-Overlapping | Total<br>Training<br>set size | Testing<br>set size |
| --- | --- | --- | --- | --- | --- | --- | --- |
| DS1 | 282 | 10 | 50 | 141 | 14:15 | 155:156 | 126:127 |
|  | 282 | 10 | 60 | 169 | 11:12 | 179:180 | 102:103 |
|  | 282 | 10 | 70 | 197 | 8:9 | 205:206 | 76:77 |
|  | 282 | 10 | 80 | 225 | 5:6 | 230:231 | 52:51 |
|  | 282 | 10 | 90 | 253 | 2:3 | 255:256 | 26:27 |
|  | 282 | 9 | 50 | 141 | 15:16 | 156:157 | 125:126 |
|  | 282 | 9 | 60 | 169 | 12:13 | 181:182 | 101:102 |
|  | 282 | 9 | 70 | 197 | 9:10 | 206:207 | 75:76 |
|  | 282 | 9 | 80 | 225 | 6:7 | 231:30 | 50:51 |
|  | 282 | 9 | 90 | 253 | 3:4 | 256:257 | 25:26 |
|  | 282 | 8 | 50 | 141 | 17:18 | 156:157 | 123:124 |
|  | 282 | 8 | 60 | 169 | 14:15 | 183:184 | 99:98 |
|  | 282 | 8 | 70 | 197 | 10:11 | 207:208 | 74:75 |
|  | 282 | 8 | 80 | 225 | 7:8 | 232:233 | 49:50 |
|  | 282 | 8 | 90 | 253 | 3:4 | 256:257 | 25:26 |
|  | 282 | 7 | 50 | 141 | 20:21 | 161:162 | 120:121 |
|  | 282 | 7 | 60 | 169 | 16:17 | 185:186 | 95:96 |
|  | 282 | 7 | 70 | 197 | 12:13 | 204:205 | 77:78 |
|  | 282 | 7 | 80 | 225 | 8:9 | 233:234 | 48:49 |
|  | 282 | 7 | 90 | 253 | 4:5 | 257:258 | 24:25 |
| DS2 | 192 | 10 | 40 | 76 | 11:12 | 87:88 | 104:105 |
|  | 192 | 10 | 50 | 96 | 9:10 | 105:106 | 86:87 |
|  | 192 | 10 | 60 | 115 | 7:8 | 122:123 | 69:70 |
|  | 192 | 10 | 70 | 134 | 5:8 | 139:140 | 52:53 |
|  | 192 | 9 | 40 | 76 | 12:13 | 88:89 | 103:104 |
|  | 192 | 9 | 50 | 96 | 10:11 | 106:107 | 85:86 |
|  | 192 | 9 | 60 | 115 | 8:9 | 123:124 | 68:69 |
|  | 192 | 9 | 70 | 134 | 6:7 | 140:141 | 51:52 |
|  | 192 | 8 | 40 | 76 | 14:15 | 90:91 | 101:102 |
|  | 192 | 8 | 50 | 96 | 12 | 108:109 | 83:84 |
|  | 192 | 8 | 60 | 115 | 9:10 | 124:125 | 67:68 |
|  | 192 | 8 | 70 | 134 | 7:8 | 141:142 | 50:51 |

The symbol (:) between numbers implies when splitting the lines across the traits some traits might have the first value and others have the second value. Splitting of lines across the traits was repeated 50 times. DS1 includes 282 NDSU pea lines while DS2 includes 192 USDA accessions.

**Supplementary Figure 1:**
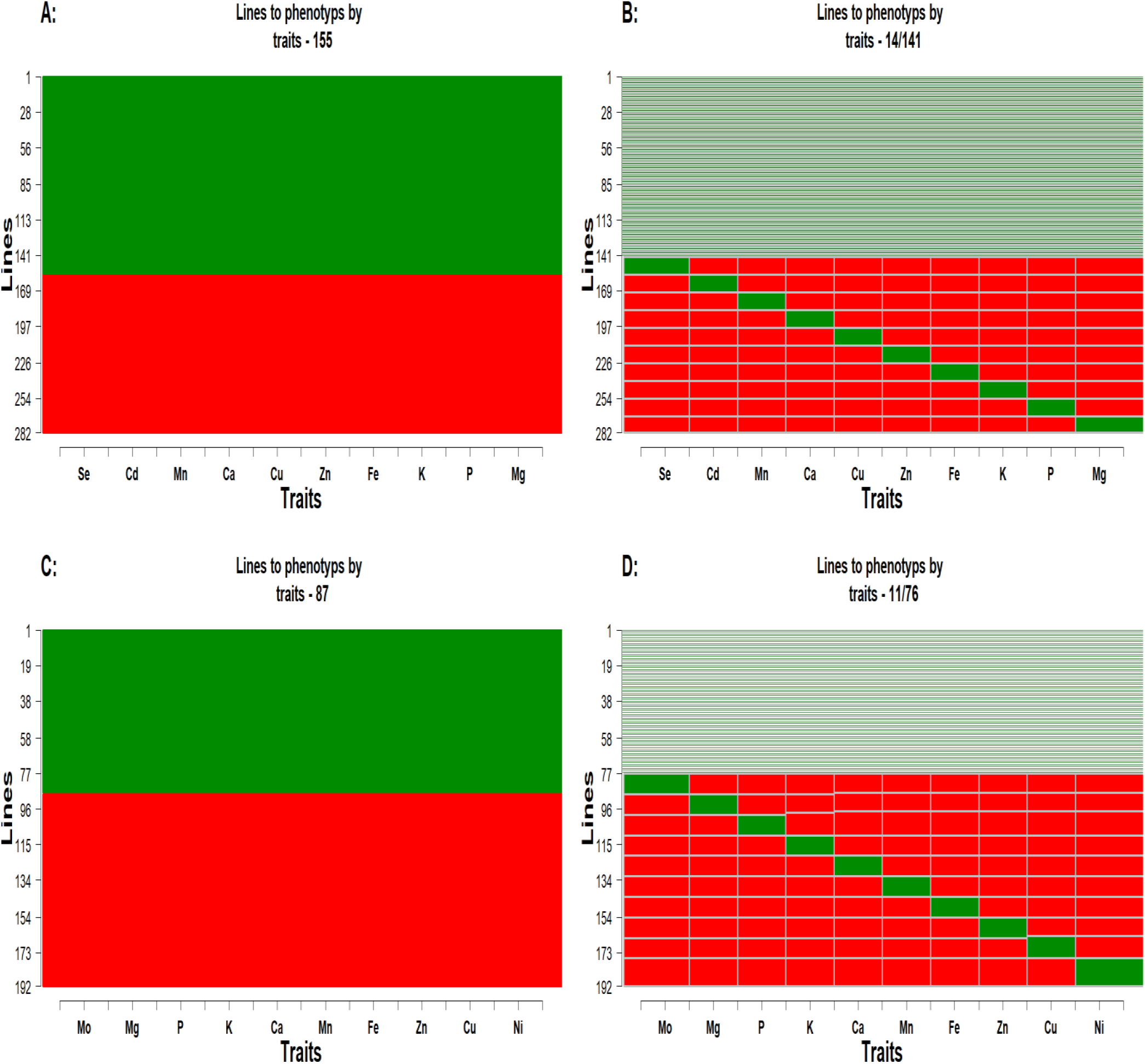
Allocation of lines to 10 traits in DS1 (A and B) and DS2 (C and D) respectively. In A and C, the green section corresponds to 155 and 87 lines in DS1 and DS2 with phenotypic data for all traits using partially balanced phenotyping. While the red implies un-phenotyped lines across traits in which the genetic value will be predicted. The B and D are sparse phenotyping strategy, each column represents a trait and the green sections correspond to 14 and 11 lines in DS1 and DS2 unique to each trait while the light green section denotes 141 and 76 lines (Approx. 50% of 282 lines in DS1 and 40% 192 lines in DS2) common to all traits.

